# Spatial analysis of ligand-receptor interaction in skin cancer at genome-wide and single-cell resolution

**DOI:** 10.1101/2020.09.10.290833

**Authors:** M Tran, S Yoon, M Teoh, S Andersen, PY Lam, P Purdue, A Raghubar, SJ Hanson, K Devitt, K Jones, S Walters, ZK Tuong, A Kulasinghe, J Monkman, HP Soyer, I Frazer, Q Nguyen

## Abstract

The ability to study cancer-immune cell communication across the whole tumor section without tissue dissociation is needed for cancer immunotherapies, to understand molecular mechanisms and to discover potential druggable targets. In this work, we developed a powerful experimental and analytical toolbox to enable genome-wide scale discovery and targeted validation of cellular communication. We assessed the utilities of five sequencing and imaging technologies to study cancer tissue, including single-cell RNA sequencing and Spatial Transcriptomic (measuring over >20,000 genes), RNA In Situ Hybridization (multiplex 4-12 genes), digital droplet PCR, and Opal multiplex protein staining (4-9 proteins). To spatially integrate multimodal data, we developed a computational method called STRISH that can automatically scan across the whole tissue section for local expression of gene and/or protein markers to recapitulate an interaction landscape across the whole tissue. We evaluated the unique ability of this toolbox to discover and validate cell-cell interaction *in situ* through in-depth analysis of two types of cancer, basal cell carcinoma and squamous cell carcinoma, which account for over 70% of cancer cases. We expect that the approach described here will be widely applied to discover and validate ligand receptor interaction in different types of solid cancer tumors.

## Introduction

Cell-to-cell communication underscores a dynamic cellular ecosystem that develops, evolves, and responds to environmental factors. The implications and roles of cell-to-cell communication have been investigated extensively, particularly in cancer, using a wide range of *in vitro* and *in vivo* techniques, albeit at different scales and resolutions (Brücher et al., 2014). Breakthroughs arising from discoveries in cell communication have led to important clinical applications. A classic example is the interaction via immune checkpoint proteins (Pardoll, 2012). Tumor cells, tumor infiltrating lymphocytes and tumor associated myeloid cells express inhibitory PD-L1/CTLA4 ligands to engage PD-1 receptors on cytotoxic T cells and CD80/86 receptors on myeloid cells, effectively blocking immune activation against the tumor cells. The discovery has led to applications of using monoclonal antibodies that specifically target this ligand-receptor (L-R) interaction as a form of immunotherapy, allowing immune cells to suppress cancer growth (Weiner et al., 2012). Therapies targeting these two pairs of ligand-receptors have transformed the management of several cancers, including melanoma, renal cell carcinoma, bladder cancer, head and neck cancer, and many others (Ott et al., 2017). Notably, often less than 20% of patients respond to a single immunotherapy, including common cancer types like breast, colon and prostate cancer (Ott et al., 2017), and hence the urgent need to combine therapies, for example by using antibodies against both PD-L1, CTLA-4 and/or PD-1 (Ott et al., 2017). However, mechanisms of action for combinational immunotherapies remain elusive (Wei et al., 2019) and the number of druggable targets for cancer-immune cell interaction is extremely limited. Therefore, research to explore and advance understanding of known and new ligand-receptor pairs in the context of tumor-immune cell interaction within a tumor is extremely important for the further development of immunotherapies (Weiner et al., 2012; Helmy et al., 2013).

Most ligand-receptor (L-R) interaction research so far has focused on the use of fluorescently-conjugated antibody-based methods, that are relatively low-throughput due to only being able to assess protein levels of a few target molecules and results are often based on a small number of cells at a time. Analysis of the transcriptome provides a means towards high-throughput L-R screening assays; however, a comprehensive and unbiased pipeline from discovery to validation of cellular communication from transcriptomics data within an intact tissue is still lacking (Browaeys et al., 2020; Efremova et al., 2020). In this work, we aim to establish a pipeline to study L-R interaction of cancer and immune cells across the whole tissue section, using skin cancer as a model. Recent developments in single-cell RNA-seq (scRNA-seq) have provided the opportunity to dissect gene expression profiles at single cell level. While inference methods to predict cell interaction using single-cell data are being developed (Browaeys et al., 2020; Efremova et al., 2020), they still lack the ability to infer spatial context of the interaction. Spatial transcriptomics (ST-seq) and RNA *in situ* hybridization (ISH) technologies overcome these limits and enable the study of (target) gene expression in undissociated tissue sections, maintaining tissue integrity (Salmén et al., 2018).

In this work, we assessed the utility of combining five complementary technologies to study and validate L-R interaction between immune and cancer cells in in skin cancer tissue. These techniques include scRNA-seq, ST-seq, RNA-ISH, droplet digital PCR (ddPCR), and Opal multiplex immune histochemistry (IHC) assay.

ST-seq measures barcoded gene expression in spots printed onto a functional glass slide (Salmén et al., 2018), which captures mRNA released from a tissue section, preserving the cell morphology. ST-seq has been applied to study the gene expression landscape of tissues and diseases, such as prostate cancer (Berglund et al., 2018; Ji et al., 2020), pancreatic cancer (Moncada et al.), melanoma (Thrane et al., 2018). However, ST-seq still has not achieved single-cell resolution per spatial spots (1-9 cells/spot), and the number of cells as well as the transcriptome quality that can be captured in each spot depend on the tissue context. These shortcomings of ST-seq can be overcome by a targeted RNA-ISH approach to visualize the cell interaction through detecting L-R at a single cell level. The RNAscope HiPlex assay (ACD Bio) has been developed based on the RNA-ISH technique and improved on the signal amplification and background suppression process compared to the previous version, allowing for visualization and detection of mRNA at near single molecule sensitivity. Using two adjoining spacers up to 50 base regions, the probes (spacers) specifically bind to target gene sequences, and then a pre-amplifier binds on the upper region of the spacer, maximizing fluorescence intensity. The technology allows researchers to simultaneously detect up to 12 single target genes on the same tissue section through fluorophore cleavage steps. In addition, ddPCR is another powerful method that can be used to confirm the expression of target genes in the same cancer tissue block used for RNAscope and ST-seq, achieving extremely high sensitivity and accuracy. ddPCR is the third generation of the PCR assay and does not require a standard curve for absolute quantification of target genes (Kuypers and Jerome, 2017; Li et al., 2018). This technique generates millions of droplets that each contains an input molecule and then partition them to be read as either a positive or negative signal according to fluorescence amplitude that is presented. Extending from measuring RNA, we implemented another technique to detect protein, covering the whole tissue and at subcellular resolution. Opal multiplex IHC can measure 4-7 proteins on the same tissue.

We combine the five experimental techniques into a powerful toolbox (STRISH) designed for cell-cell interaction analysis, uncovering the interaction of immune cells and cancerous cells in the whole tissue section. The combination enabled us to build an experimental and analytical pipeline to comprehensively discover and validate L-R interaction at the transcriptional level. To identify L-R pairs at a transcriptome-wide scale, we applied two approaches, starting with scRNA-seq and followed by ST-seq, using skin cancer samples. Most ligands and receptors are expressed at a relatively low level, leading to a high possibility for the under-detection of those genes by using scRNA-seq and ST-seq. Therefore, we applied scRNA-seq and ST-seq at the discovery phase to detect potential L-R pairs, followed by the sensitive validation of a selected set of genes, by using single-molecule detection methods: RNAscope and ddPCR. RNAscope generates spatial information at single-cell resolution and ddPCR is approximately 1,000 times more sensitive than RNA-seq. We then applied Opal IHC to validate the interaction at protein level.

Together, the five technologies allow us to perform end-to-end L-R analysis, from unbiased discovery with ST-seq and scRNA-seq to targeted validation at RNA and protein levels by RNAscope, ddPCR and OPAL multiplex IHC. Importantly, we developed an accompanied computational pipeline to process these multimodal datatypes (STRISH). We demonstrated the utilities of the pipeline through in-depth analysis of two types of the most common skin cancer, covering three pairs of L-R ranging from low to high abundance. These pairs include interleukin-34 (IL34), colony-stimulating factor 1 receptor (CSF1R), THY1 (also known as CD90), Integrin subunit alpha M (ITGAM, also known as CD11b), PD-1 and PD-L1. Our work provided the important assessment of the genomics and imaging technologies that can be used to discover, validate and understand immune-cancer cell interaction within a tumour section, commonly used in histopathological assessment of cancer.

## Results

### Genome-wide analysis of ligand-receptor interaction using single-cell RNA sequencing

scRNA-seq and spatial transcriptomics are the five technologies, as shown in Figure 1A, that allow for unsupervised prediction of cell-cell interaction. To infer potential L-R pairs that are likely used as means of intercellular communication, we applied common inference methods, CellPhoneDB (Efremova et al., 2020) and NicheNet pipeline (Browaeys et al., 2020) on our scRNA-seq dataset of matched samples, containing a squamous cell carcinoma sample tissue (SCC) and a normal sample tissue from the same patient (Figure 1G, H; Figure S1A). NicheNet combines expression data with prior knowledge on gene signaling and gene regulatory networks to predict L-R pairs used by interacting cells (sender and receiver cells). If the inference approach using scRNA-seq was sufficiently sensitive, we expect to detect known IL34-CSF1R and THY1-ITGAM interaction. The two pairs were not among the significantly detected pairs predicted by NicheNet (**Supplemental Figure S1A**). To examine the relatedness of cells coexpressing the L-R pairs, we visualise L-R expression relative to cell cluster information in the UMAP space (**Supplemental Figure S1B-D**). We found evidence of cells in the same clusters that expressed IL34 and CSF1R (**Supplemental Figure S1C**). However, THY1 and ITGAM were only expressed by a few cells with no distinct patterns of coexpressio (**Supplemental Figure S1D**). The results from examining THY1-ITGAM suggested the insensitivity of using scRNA-seq alone to study cell communication. Overall, we found that scRNA-seq can be utilized to study cell communication but is limited by insensitivity and a lack of spatial information. Therefore, we next assessed spatial transcriptomics to detect L-R interaction by integrating spatial information.

**Figure 1.**
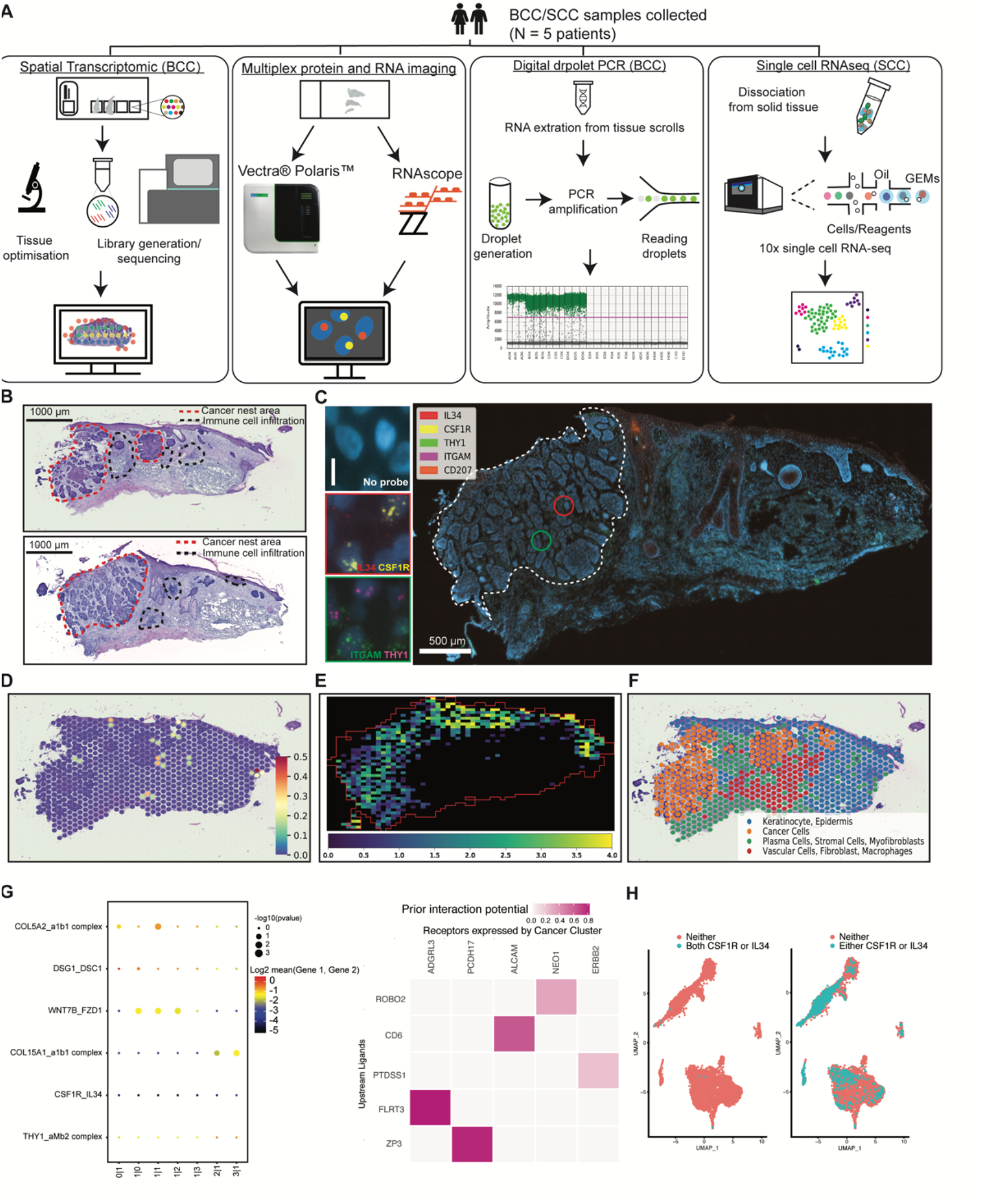
A technological and computational pipeline to study interaction across the whole tissue section. **(A)** A workflow illustrating the combination of five technologies to study L-R interaction in skin cancer tissue. The technologies include Visium Spatial Transcriptomic (ST-seq), RNAscope Hiplex, Opal multiplex protein assay, automated ddPCR, and scRNA-seq. **(B)** The annotated H&E images of the two adjacent tissue sections that were used for Visium ST analysis (top) and the consecutive section of the same block used in RNAscope assay (bottom). **(C)** Target RNA molecule expression at a single cell level using RNAscope assay and the visualization of the local co-expression of two pairs L-R including IL34-CSF1R and THY1-ITGAM (left panel). **(D)** Using ST-seq analysis to determine local co-expression level of IL34-CSF1R. **(E)** STRISH pipeline to detect the local co-expression of IL34-CSF1R using RNAscope imaging data. The red solid line boxes indicate the regions where they are consistent in ST-seq and RNAscope analysis. **(F)** Spatial feature plots of the four distinct clusters defined by Louvain graph-based clustering. The annotation was based on differential gene and pathway analysis. The distribution of the cluster annotated as cancer was consistent with the location of cancer nests in upper image of **(B). (G)** The inference of cell-cell interaction through ligand-receptor using CellPhoneDB (left) and NicheNet (right) pipelines using ST-seq data. For CellPhoneDB prediction, top four highly active pairs of ligand-receptor and the two target pairs were selected for visualization. **(H)** The UMAP feature plots highlighted the cells that expressed both CSF1R and IL34 (Cyan dots, left plot) and either CSF1R or IL34 (Cyan dots, right plot).

### Genome-wide analysis of ligand-receptor interaction using spatial transcriptomics

We postulated that cell-cell communication could be better assessed by using ST-seq, which preserves neighborhood information of interacting cells. We performed ST-seq on one tissue section from a BCC patient (Figure B, D, F). Adjacent tissue section was used for RNAscope analysis as discussed later (Figure B,C,E). We annotated the cancer and immune cells in the two hematoxylin and eosin (H&E) images for adjacent tissue sections for RNAscope and for ST-seq. The major cell types include cancer nests, immune infiltrations and non-cancer regions (**Figur 1B**). From the same block, one section was used for ST-seq and another for RNAscope, top and bottom sections respectively (**Figure 1B**). The ST-seq and RNAscope data from these two sections were subsequently used to model cell-cell communication and the results were compared. The consistency in predicted interaction activities allows us to validate the results from ST-seq using RNAscope data (**Figure 1C-E; Supplementary Figure 2A**). The modelling and detailed comparisons will be discussed in the later section.

With ST-seq approach, we reasoned that each spot in the Visium slide contains a mixture of cells and these cells can communicate if both ligand and receptor are detected in the spot. In addition, interaction between cells from two neighboring spots, through paracrine signaling, can likely occur if these spots display L-R co-expression. Therefore, local co-expression of L-R pairs within a spot or between neighboring spots suggests possible interaction. We generated the cell-cell communication activity map across the whole tissue for IL34-CSF1R from ST-seq data (**Figure 1D**) using our cell-cell interaction algorithm implemented in stLearn software based on the above reasoning (Pham et al.). The heatmap of interaction in the ST-seq slide showed the high intensity of cell interaction within and surrounding the tumor areas as well as the immune infiltration regions. Further supporting evidence for IL34-CSF1R interaction was obtained from the pathological annotation (Figure 1B) and data-driven functional annotation of cell types. Unbiased clustering of gene expression using a graph-based approach on the 750 spots in the tissue section resulted in the demarcation of the 4 distinct groups (**Figure 1F**). The cell types were annotated by running differentiation gene expression (**Supplementary Figure S2E**) followed by gene ontology enrichment analysis. We identified two major populations in cluster 1, observed in normal skin including keratinocyte (expressing CSTA, KRT6A), epidermis (expressing KRT10, KRT14). Meanwhile other 3 clusters expressed markers for cancer (SFRP5, GLI1), stromal cells (CD40, THY1), and macrophages (CXCL12, CD37). The distribution of the 4 cell types overlapped well to the skin morphology and cancerous areas defined in H&E images. The interaction activities were highest in the area with more cancer and immune infiltration cells, particularly in the epidermal compartment.

Interestingly, we also found that computational methods that did not use spatial information failed to detect the expected L-R interaction. From the CellPhoneDB analysis pipeline (Efremova et al., 2020), a total 1024 possible combinations of L-R were tested using the ST-seq dataset. The dot-plot in **Figure 1G** demonstrated the low significance of the two L-R pairs, IL34-CSF1R and THY1-amb2 complex (ITGAM pathway) in comparison to the 4 other CellPhoneDB highest significance of the interaction pairs. We further applied NicheNet (Browaeys et al., 2020) workflow on the ST-seq data to integrate prior knowledge into the prediction. While NicheNet could identify interaction pairs that were directly linked to cancer (i.e., CD6-ALCAM), the prediction disregarded the spatial information and failed to find possible interaction. The expression plots of the four target markers overlayed to the tissue section (**Supplementary Figure S2B, S2C**) illustrated the low abundance across the tissue. The low depth reads and the inherent dropout (randomly misdetecting molecules due to the sequencing protocol) of ST-seq (and scRNA-seq). Therefore, for the noisy (high dropout) data, computational methods without spatial information like CellPhoneDB and NicheNet likely miss-identify meaningful cell-cell communication (**Figure 1 G,H**), while spatial interaction analysis method, like stLearn, could detect such signalling events (**Figure 1D**). However, independent validation experiments are required. Next, we describe RNAscope and our novel STRISH pipeline as a powerful experimental method for validating cell-cell interaction within spatial tissue sections.

### STRISH, a computational pipeline to map ligand-receptor interaction activities across the whole tissue (interaction landscape) using RNAscope data

To overcome the limitations in detection sensitivity due to a lack of signal in the cancer region from the sequencing methods (scRNA-seq and ST-seq) and to achieve single cell resolution, we implemented RNAscope HiPlex assay. The whole fluorescent microscopy images of the tissue sections were captured at 40x magnification to determine for RNA interaction at cellular resolution. Three different fluorophores (Cy3, Cy5 and Cy7) were used in two iterative wash-stain rounds to label the five distinct target genes. Following image acquisition, the first image for THY1, IL34, and CSF1R and the second image for CD207 and ITGAM were overlaid to generate a single merged image, for analysis of L-R interaction (**Figure 1C**). The zoom-in images from a cancer nest area (marked as a white dashed line, and two red/green circles - IL34 and CSF1R in a red box and THY1 and ITGAM in a green box) show distinct cell to cell interaction for each of pair compared to no signal in the ‘Negative’ control.

To automate and improve the accuracy of detecting L-R interaction from fluorescent data, like RNAscope, we developed a computational pipeline STRISH. The STRISH pipeline consists of two phases, starting with a cell local co-expression detection step to define spatial neighbourhood, followed by scoring and visualizing the local co-expression level (**Figure 2A, Method Algorithms 1**,**2**). Local co-expression is defined as the expression within a tissue area containing fewer than a threshold number of cells (depending on tissue types), for example fewer than 100 cells. The pipeline runs a series of positive-cell detection iterations to find regions that contain lower than the predefined threshold of neighboring cells and subsequently determines the number of L-R co-expression within these regions (refer to the Method section about STRISH algorithm). Across the tissue samples, we observed considerable interaction of IL34-CSF1R around the areas where the cancer nests are located in both BCC and SCC, particularly in the epidermal compartments (**Figure 1E, 2B; Supplemental Figure S3A, S3B**). Interestingly, compared to RNAscope data we observed a similar pattern in ST-seq data, with many spots that were predicted to have cell-cell communication through IL34-CSF1R located in cancer and epidermis regions.

**Figure 2.**
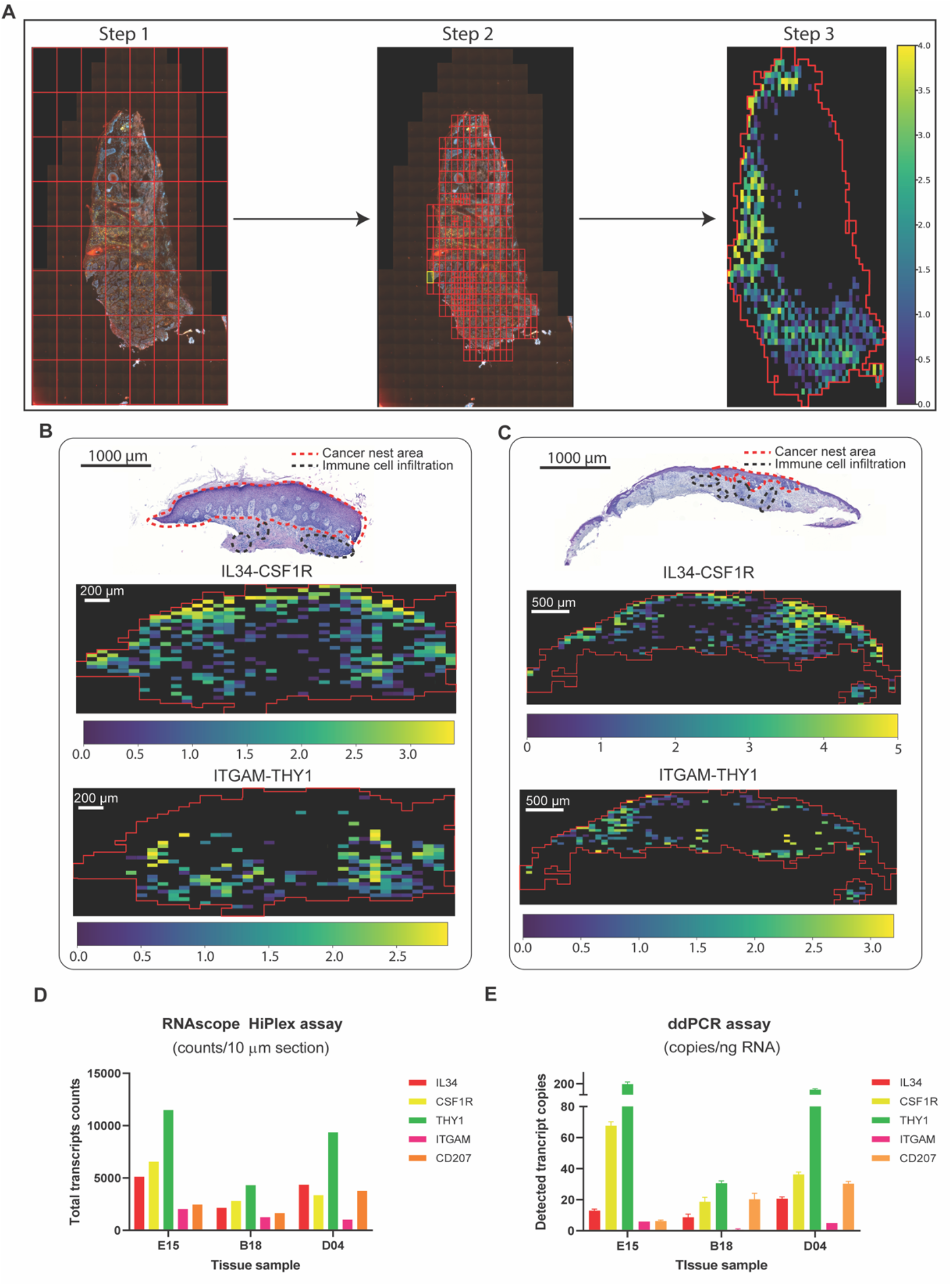
Detection of target RNAs in skin cancer patients by collective transcriptomic and genomic methods. **(A)** STRISH analysis steps from raw RNAscope data to create a tissue-scale heatmap (activity map) of local co-expression of target mRNAs. **(B)** and **(C)** From top to bottom: annotated H&E images and corresponding heatmaps (activity maps) of local co-expression of two BCC patient samples, ID-B18 and ID-D04. The heatmaps of the co-expression of the two L-R pairs IL34-CSF1R and THY1-ITGAM of the two patients are shown in the middle and the bottom rows, respectively. **(D)** and **(E)** The absolute copy number of the target mRNAs counted from the two assays.

**Algorithm 1:**
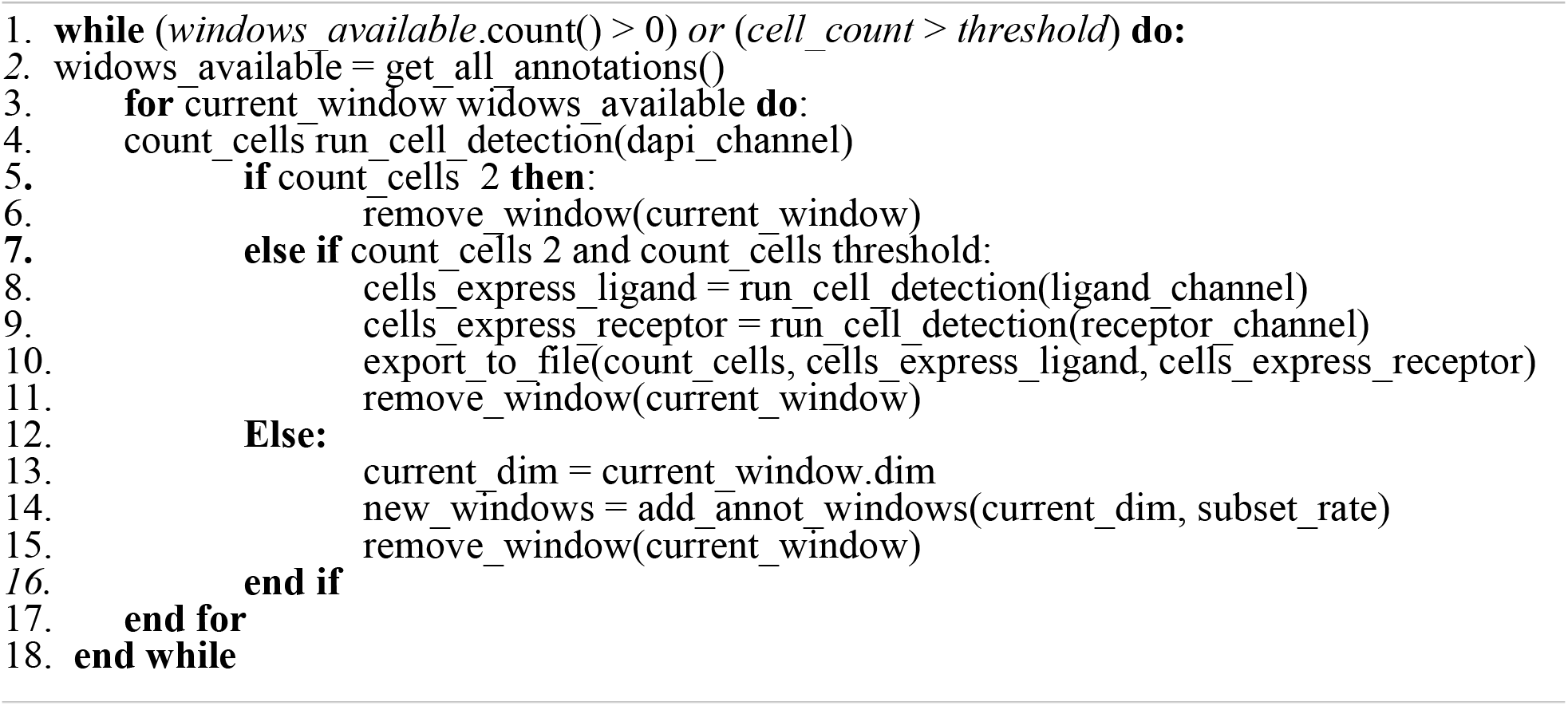
Cell detection by window scanning

**Algorithm 2:**
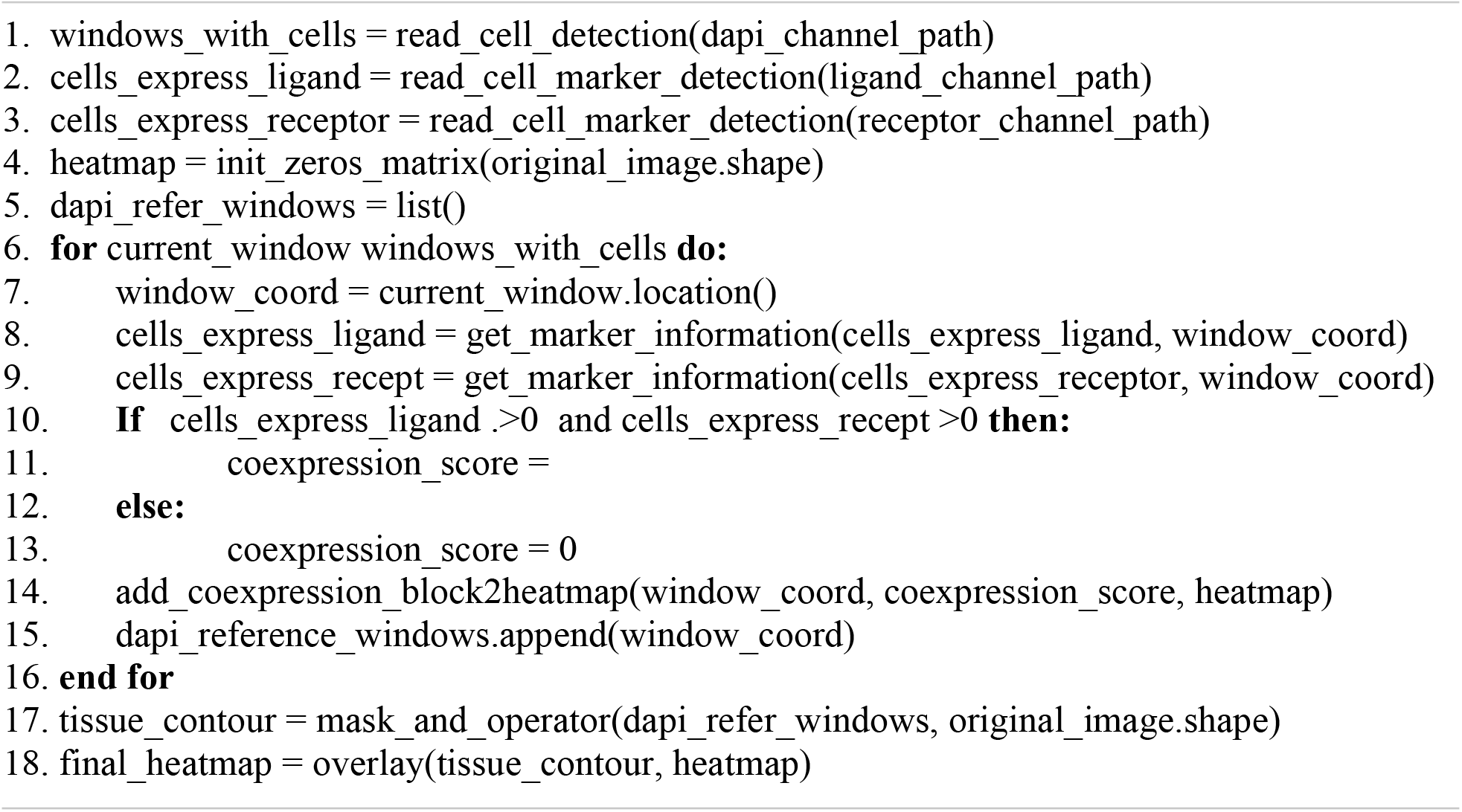
Quantifying local co-expression and visualization

We further assessed the performance of STRISH by measuring the interaction of another L-R pair, THY1-ITGAM. We found lower signal and less wide-spread interaction of THY1-ITGAM that were localized to the dermal compartments (**Figure 2B; Supplemental Figure S3A, S3B**). By comparing the number of the tissue regions (through STRISH windows) where local co-expression was found, we noticed the average count of the window for THY1 and ITGAM to local co-expression was 2.5 times less than the one for the pair of IL34-CSF1R. Similarly, we calculated the values of the normalized local co-expression in the same area per STRISH window for IL34 were higher than that of ITGAM. We noted that most of the STRISH-detected local co-expression of THY1 and ITGAM were clustered densely around the adjacent areas of the immune cell infiltration. Notably, STRISH can quantitate the level of interaction by counting local-expression events, as shown in the bar graph (**Figure 2D, formula (2)**). Overall, we found that STRISH can detect cell-cell communications more sensitively than ST-seq and scRNA-seq.

### Confirmation of ligand-receptor expression by ddPCR

While RNAscope is expected to be able to detect single molecules in each cell, scanning through a whole tissue section with millions of cells may lead to noise and reduced accuracy due to tissue heterogeneity. ddPCR, on the other hand, enables to sensitively detect and quantify single molecules from tissues, albeit spatial information is absent. To further confirm the presence of ligand and receptor in the tissue, we performed ddPCR on the same cancer tissue block (**Supplementary Figure S4**). The transcript copy number per input RNA of each gene was presented in a bar plot (**Figure 2E**). ddPCR detected both L-R pairs and interestingly the result from ddPCR was highly consistent to that from. We used Pearson’s correlations to compare the quantitative analysis using RNAscope and ddPCR (**Figure 2D, 2E**) and found correlation values between the two assays as 0.95, 0.89 and 0.94 for the patient samples tested, ID-E15, B18 and D04, respectively. The ddPCR results support the quantitativeness of using RNAscope for measuring target gene expression, suggesting the suitability of using RNAscope for detecting and quantifying cell-cell interaction.

### Extending the application of STRISH from RNA-ISH to detect L-R interaction using protein fluorescence data

We generated multiplex protein immunofluorescence data for a SCC cancer sample using multispectral imaging with primary antibodies-Opal pairing technologies (Akoya biosciences). On the same tissue section, seven cancer and immune markers were detected simultaneously, including a L-R pair (PD-1 and PD-L1), PanCK (epithelial cancer marker), CD8 (T cell marker), CD68 (Macrophage marker), FoxP3 (T regulatory cell marker), (**Figure 3A**). We implemented the STRISH pipeline to detect the local expression levels of the L-R pair PD-1 and PD-L1, which are well known to be signalling molecules for immune and cancer cell interaction (Pardoll, 2012), (**Figure 3 B**). We divided the fluorescent tissue image into multiple smaller areas containing fewer than 100 cells, defined as a local neighbourhood region (refer to Algorithm 1, Method Section). We then quantified the cell expression on the window for either PD-1 or PD-L1 (refer to Algorithm 2, Method Section). The activity map of PD-1 and PD-L1 interaction is shown as a tissue heatmap with colors representing the normalised count (normalisation of all cells within each window) of PD-1 and PD-L1. The automated STRISH detected regions with high co-expression of PD-1 and PD-L1 were visually examined, separately for each of the two channels, PD-1 and PD-L1 (**Figure 3C**). Similar to the RNAscope data, we showed that STRISH could detect PD-1 and PD-L1 local co-expression accurately. Visual inspection of positive and negative windows (**Figure 3C**) shows the unique new feature of STRISH in that it detects local co-expressing cells of the L-R pairs, in contrast to other methods that only detect the co-expression of two or more proteins from the same cell.

**Figure 3.**
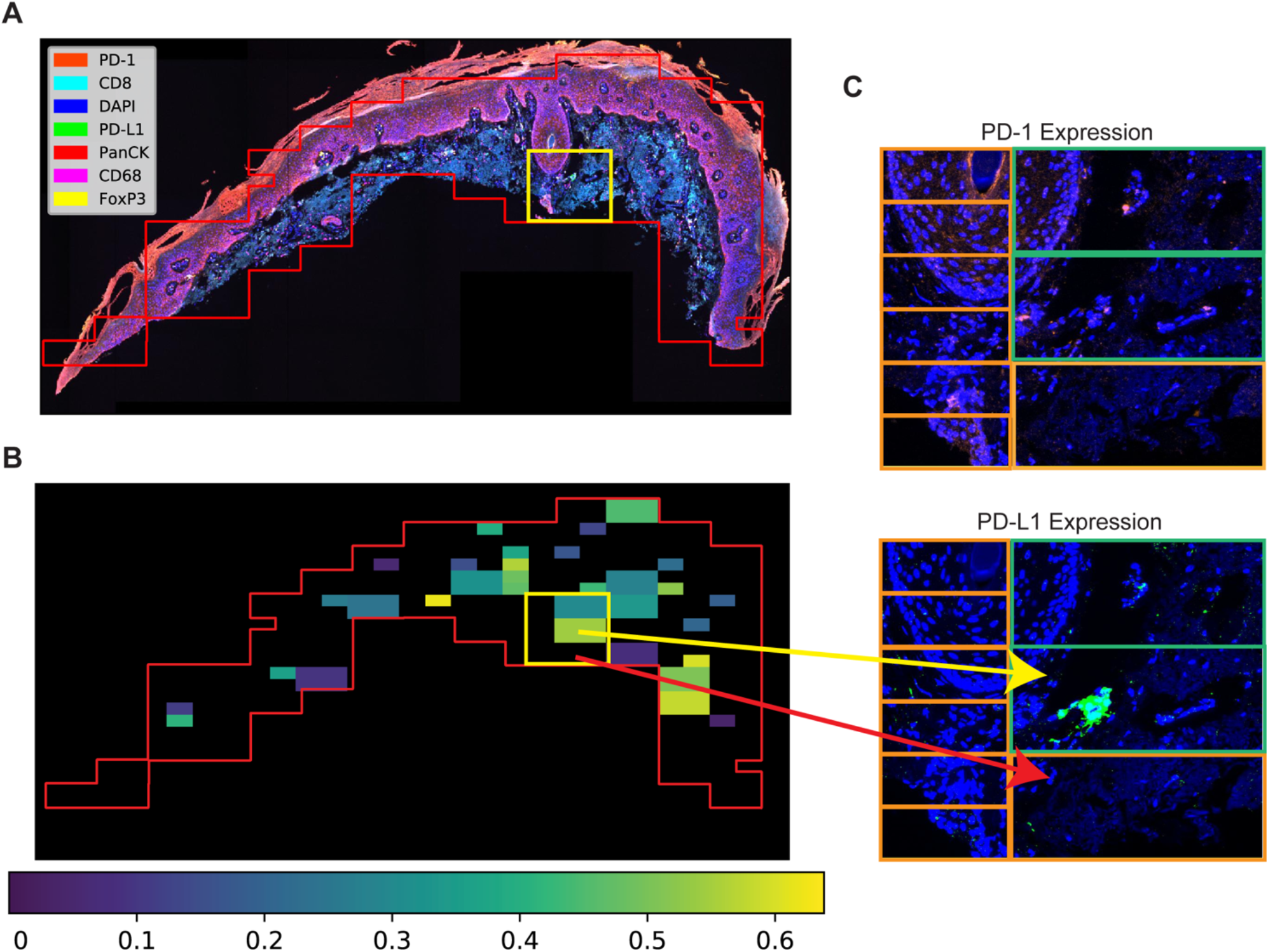
STRISH application to protein data. **(A)** A multispectral image captured using MOTiF^™^ PD-1/PD-L1 panel of an SCC cancer tissue section. **(B)** STRISH heatmap (activity map) suggesting the tissue locations with high (yellow color) or low (dark color) level of protein local co-expression for the ligand-receptor pair PD-1 and PD-L1. The tissue contour was plotted using the windows of neighboring cells identified by STRISH. **(C)** A close-up visualization of the areas identified as the existing cells local co-expression of PD-1 and PD-L1 by STRISH. The windows that STRISH identified the local co-expression were highlighted in green windows while the orange windows showed the control regions where there is no co-expression detected.

## Discussion

We presented a technological and computational end-to-end pipeline from discovery to validation of L-R interaction across the whole spatial landscape of a tumour tissue section. We assembled and evaluated 5 technologies used for studying L-R interaction. We generated multimodal data using these 5 methods and examined over 1000 L-R pairs in scRNA-seq and ST-seq data, followed by in-depth analyses of three L-R pairs in two types of two non-melanoma skin cancers, which account for about 70% of all cancer cases (Garcovich et al., 2017). The pipeline allows us to quantitatively and visually assess the expression of target transcripts while maintaining the physiological spatial information in undissociated tissue sections. Computationally, we developed and demonstrated the utility of an imaging analysis pipeline called STRISH to automatically detect tissue regions with high cell-to-cell interaction activities using RNAscope RNA and Opal multiplex protein images. The STRISH results are important orthogonal validation of the predictive results from scRNA-seq and ST-seq analysis.

STRISH is a powerful computational pipeline to quantitatively characterize cell-cell interaction by automatic scanning of a local expression of a target gene in a RNAscope image to recapitulate an interaction landscape across the whole tissue. The pipeline also consists of a robust image registration step to merge two separate images from multiple imaging rounds, with potential extension to merging two adjacent tissue section. This utility allows for data integration, increasing the multiplex capacity to detect more L-R in one tissue section. Furthermore, STRISH effectively detects cells using non-overlapped windows followed by computationally evaluating the level of local L-R co-expression in each window. Existing image analysis tools have been developed to search for colocalization of two gene/protein markers within each individual cell (Bolte and Cordelières, 2006). STRISH, in contrast, finds co-expression between neighbouring cells rather than within one cell, thereby identifying potential paracrine interaction between cells that are in proximity to each other. The number of neighbouring cells can be determined by users, allowing the flexibility to different biological systems, such as cancer types. Eventually, the interaction activity can be visualized in a heatmap covering the whole tissue landscape. STRISH, therefore, allows us to model L-R autocrine and paracrine interaction landscape across the whole tissue section. For the generalisation of the method, we assessed the local co-expression of IL34-CSF1R and THY1-ITGAM using STRISH for RNA-ISH multiplex assay and PD-1 vs PD-L1 for Opal protein assay. Importantly, the extension of STRISH application to immunofluorescence data suggests the very broad applicability of this analysis pipeline to the vast amount of protein fluorescence imaging data.

Our detection of CSF1R and CSF1/IL34 interaction between cancer and immune infiltrating cells at the epidermal layers was consistent to the biological context of the skin cancer. The signaling interaction between CSF1R and CSF1/IL34 is well known to regulate macrophage differentiation (Lin et al., 2008). IL34, CSF1 and their receptors co-expressed within the immune cell infiltrated area and regulated the different downstream signaling pathways in breast cancer (Zins et al., 2018). CSF1R is known to be expressed on CD1a/CD207 Langerhans cells in human epidermis and stratified epithelial (Lonardi et al., 2020), where it can be found in high abundance in the basal and squamous layers of the epidermis, and is involved in anti-tumoral immune responses (Pogorzelska-Dyrbuś and Szepietowski, 2020). Consistent with the previous findings, we observed that the co-expression of IL34-CSF1R L-R pair is high in/near cancer nest area in all the patient samples analyzed.

Similarly, we found that while the co-expression of ITGAM-THY1 was not as high as the one of IL34-CSF1R with respect to tissue morphology related to cancerous areas, it was correlated with regions of infiltrated immune cells rather than cancer nests. This is concordant with the literature, suggesting that this L-R pair may be involved in migration of leukocytes to damaged cells and initiate host defence (Wetzel et al., 2004). THY1 is one of the cell surface markers for T-lymphocytes and mesenchymal stromal cells (Uder et al., 2018). The interaction of THY1 and ITGAM, an integrin molecule marking myeloid cell populations such as monocytes and polymorphonuclear granulocytes, has been reported (Wetzel et al., 2004). The interaction is shown to be involved in leukocyte migration in injured or inflammatory tissue, leading to an initiation of the function of leukocytes in host defences (Leyton et al., 2001).

Our results suggest RNAscope Hiplex assay to be quantitative, sensitive and high-resolution in detecting L-R interactions. RNAscope detection is more suitable than IHC in precancerous dysplastic nodules, helping early monitoring and diagnosis for patients with high risk of diseases (Bakheet et al., 2020). Additionally, our results from the absolute quantification of the target genes from ddPCR analysis (using adjacent tissue sections of the same tissue blocks) suggests that the RNAscope is able to accurately measure the expression of L-R pairs at the tissue level. This observation is important because RNAscope assay quantifies from a microscopic image that captures the fluorescent transcripts, which usually semi-quantitative. The high correlation to ddPCR results suggests the high accuracy of RNAscope assay. ddPCR measures the total target transcript number in the input RNA sample, detecting RNA as sensitive as at a single molecule resolution. The combination of RNAscope and ddPCR can be considered as a targeted validation approach for the unbiased, but less sensitive approach by spatial transcriptomics. The further Opal multiplex protein assay, on the other hand, validate the expression at the protein level.

Here, we propose a complete pipeline from bench work to bioinformatic analysis to study cell interaction through L-R pairs in cancer tissues. The workflow demonstrates the feasibility to discover new L-R pairs by genome-wide approaches (scRNA-seq and ST-seq), which are less sensitive but cover all genes, followed by targeted validation by high-resolution, sensitive RNAscope imaging, ddPCR, and Opal multiplex protein assays. The pipeline integrates multimodal data to assess cell-to-cell interaction in tumors using collective methodologies on a same cancer patient specimen. The combination of different technologies overcomes the inherent limitations of any individual method. Our technological and analytical platform that allow for the discovery and detailed-analyses of more than one pairs of L-R interaction in cancer tissues provides a powerful approach on finding new targets and/or understanding the mechanisms underlying options for combinatorial immunotherapies.

## Methods

### Single cell RNA analysis

Fresh shaved suspected SCC and BCC lesions and a 4mm punch biopsy of non-sun exposed skin from the same patient were collected in DMEM for immediate tissue dissociation. The tissue sections were rinsed in PBS and incubated in 8 mg/mL Dispase at 37°C for 1h, and were minced before a 3 minute incubation in 0.25% Trypsin at 37°C. To collect single cells, the suspension was filtered through a 70µm cell strainer. Cells were collected in PBS containing Fetal Calf Serum. The 10x Genomics Chromium scRNA-sequencing followed the manufacturer’s instructions, using the Single Cell 3’ Library, Gel Bead and Multiplex Kit (version 2, PN-120233; 10x Genomics). Cell numbers in each reaction were optimized to capture approximately 3,000 cells. The single-cell transcriptome libraries were sequenced on an Illumina NextSeq500, using a 150-cycle High Output reagent kit (NextSeq500/550 version 2, FC-404-2002; Illumina) as follows: 98 bp (read 2), 8 bp (I7 index), and 26 bp (read 1 - cell barcodes and UMI).

The BCL file was converted to a FASTQ file using bcl2fastq/2.17. We used CellRanger/3.0.2 for mapping to Homo_sapiens.GRCh38p10 reference. Using Seurat/3.2.0 in R/3.6.3, we removed cells with fewer than 200 or more than 5000 genes and cells with over 20% of all reads mapped to mitochondrial genes. The processed expression matrix was scaled to 10,000 reads/cell, log normalized, and only the top 2500 most variable genes were kept for PCA dimensionality reduction. The top 50 PCs were used for building a nearest neighbor graph, followed by Louvain graph-based clustering in Seurat at the resolution of 0.5 in **Supplemental Figure S1B**.

We applied NicheNet L-R prediction pipeline (Browaeys et al., 2020) on our two scRNA-seq datasets of cancer and normal samples from a patient (**Supplemental Table S1**). We ran the gene differential expression analysis for two conditions of cancer and normal. For the differentially expressed genes we used top 20 upstream ligands, then filtered the prebuilt ligand receptor network to find the corresponding upstream receptor and plotted the heatmap of potential interaction in **Supplemental Figure S1A**.

### Visium spatial transcriptomics sequencing and analysis

Tissue cryosectioned at 10µm thickness were transferred to chilled Visium Tissue Optimization Slides (3000394, 10x Genomics, USA) and Visium Spatial Gene Expression Slides (2000233, 10x Genomics, USA), and allowed to adhere by warming the back of the slide. Tissue sections were dried for one min at 37°C, fixed in chilled 100% methanol for 30 minutes and stained with hematoxylin and eosin for 5 minutes and 2 minutes as per Visium Spatial Tissue Optimization User Guide (CG000238 Rev A, 10x Genomics) or Visium Spatial Gene Expression User Guide (CG000239 Rev A, 10x Genomics). Brightfield histology images were captured using a 10x objective on an Axio Z1 slide scanner (Zeiss). Brightfield images were exported as high-resolution tiff files using Zen software. This H&E staining and imaging protocol was used to stain all skin sections for histopathological annotation in this study.

The Visium raw sequencing data in BCL format was converted to 110,782,035 FASTQ reads using bcl2fastq/2.17. The reads were trimmed by cutadapt/1.8.3 to remove sequences from poly-A tails and template-switching-oligos. We used SpaceRanger V1.0 to map FASTQ reads to the cellRanger human reference genome and gene annotation for GRCh38-3.0.0. On average, for each spot we mapped 94,710 reads and detected 1,428 genes. The count matrix of the Visium data was preprocessed to remove genes that expressed in less than three cells, followed by the normalization, log transformation and scaling. For spot clustering, we first performed gene expression normalization using spatial morphological information from the H&E image then clustered the spots using Louvain community detection algorithm. The normalization was to reduce the technical limitation in detecting lowly expressed genes. Besides, a neighborhood graph of spots was built based on the reduced dimensional space, followed by the application of Louvain community detection to group similar spots into clusters.

We performed ST-seq on one tissue section from a BCC patient (patient ID-E15). The prediction of cell-cell interaction of a pair IL34 and CSF1R analysis is produced by the stLearn package (Pham et al.) (**Figure 1C**). For L-R prediction with CellPhoneDB, we applied the default parameters (Browaeys et al., 2020; Efremova et al., 2020) and used a curated database v2.0.0. Similar to scRNA-seq analysis for cell-cell communication via L-R pairs, we applied NicheNet L-R prediction on ST-seq data. The upstream ligands were ranked by descending Pearson values and top 5 ligands were pulled out to sort out the corresponding upstream receptors.

### Multiplexed RNA in-situ hybridization with RNAscope and cell-cell interaction analysis

The following target probes to detect L-R interaction were designed by ACD probe design Team and used for the RNAscope HiPlex assay (ACD Cat. No. 324110): THY1 (ADV430611T2), IL34 (ADV313011T3), CSF1R (ADV310811T4), CD207 (ADV809521T7), and ITGAM (ADV555091T8). The assay was performed as described in the manufacturer’s user manual (ACD, 324100-UM). Briefly, a 10µm thickness tissue slide sectioned from the OCT embedded BCC or SCC tissue block was used for the assay and a consecutive section was made for a negative control. The (frozen) sections were fixed with freshly made 4% PFA for an hour followed by a dehydratio process in ethanol and then were digested with protease IV for 30 minutes at room temperature. The slide was stained with a mixture of the 5 probes to allow them to hybridize with RNAs. The negative control slide was stained with RNAscope HiPlex 12 Negative control Probe that was provided in the kit. Consequently, a specific signal was amplified with high efficiency using RNAscope HiPlex Amp 1–3 reagents. After several iterative washes using a washing buffer, the sections were then stained with RNAscope HiPlex Fluor T1–T4 reagent and were counterstained with DAPI followed by mounting with a ProLong Gold Antifade Mountant (Fisher Scientific).

The images were captured by Axio Z1 slide scanner (Zeiss) with an appropriate adjustment of each fluorescent intensity. The first round of images was performed using 4 filters including DAPI for nuclei, Cy3 for THY1, Cy5 for IL34, and Cy7 for CSF1R. For the high resolution of an image, a 40x objective was used and the Z-stack interval was set up to 1.5µm resulting in 9 of Z-slices for each slide. Completing the first round of image, the fluorophores on the slide were cleaved for the second round of imaging process. The sections were stained with RNAscope Fluoro T5 – T8 reagent and images were captured using 3 filters including DAPI for nuclei, Cy5 for CD207 and Cy7 for ITGAM. The parameters for the microscope were set up the same as the first round. The images were further processed by ZEN software (version 3.2) for manual stitching and adjusting contrast/brightness.

### STRISH computational analysis of local co-expression

To uncover the interaction of immune cells and cancerous cells in the whole BCC/SCC tissue section, we developed an analysis pipeline, called (STRISH). For detecting cell local co-expression, we utilized the built-in cell detection function from QuPath software (Bankhead et al., 2017) and employed the detection on the RNAscope stitched image produced by the Zen software (ZEISS microscopy). We developed STRISH as a python-based package for downstream analysis of cell local co-expression, as a model to predict cell-cell interaction.

During the RNAscope imaging, the slide was scanned using Z-stack interval which resulted in multiple Z-slices image. Prior to the cell detection, we applied the preprocessing step to calculate the mean signal values of all 9 of Z-slices

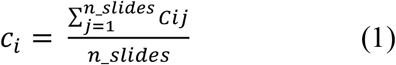

where *c*_*i*_indicates the DAPI or RNA marker channel, *j* is an iterator of the Z-slices.

To quantitatively measure the local co-expression of each window, we defined a scoring function to score the number of cells that express either ligand or receptor in the same window following the equation (2).

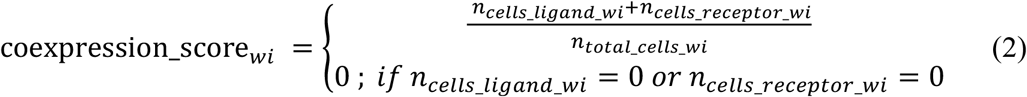

where *n*_*cells_ligand*_,*n*_*cells_receptor*,_*n*_*total_cells*_ are the number of cells that express ligand, receptor marker, and total number of cells present in the current window *wi*.

The STRISH functionality for cell local co-expression was developed to iteratively scan through the images, using a neighbourhood window detection strategy to find the target regions with cells expressing the marker of interest. First, the cell detection is initiated to cover a broad area of scanning with dimensions for width and height set to a predefined rate of the whole scan images. While iterating through the stack of all the existing windows, STRISH will discard those windows with fewer than two cells (cell count is based on DAPI signal). For each window, where the number of detected cells is greater than a threshold (user’s defined threshold depending on cancer tissue types), STRISH further splits it into smaller windows (the default rate is 50% of the current considering window dimension) and adds these smaller windows into the iteration stack (**Figure 2A**). Finally, windows that pass the cell threshold check are subjected to the next step to detect co-expression. STRISH measures each target marker expression level by applying another round of detection for markers and signal thresholding accordingly.

For the visualization of local co-expression activity, we define the threshold for positive cell with each marker by considering the number of cells in the same window to make all the windows comparable. The STRISH local co-expression level of each target pair was calculated by normalizing the total count of each pair in a window to the number of the cells detected in the same window (Formula 2). We developed a Python-based pipeline to construct the heatmap of cell local co-expression.

As there were two rounds of RNAscope imaging, we added to STRISH an image registration functionality. For the interaction of ITGAM and THY1, as the signal of the genes were captured in two separated imaging rounds with the respective stitching in post process, some variants were introduced (**Supplemental Figure S5A, S5B**). To overcome the unaligned tissue layout, we performed image registration to map one image to the other (**Supplemental Figure S5C**). The image registration is performed solely using SITK library (Lowekamp et al., 2013; Yaniv et al., 2018). Upon the registration, the same process as described above was applied to detect L-R interaction in the merged image and subsequently, the heatmap of the local co-expression was generated (i.e., **Supplemental Figure S5D**).

Our code for detecting local co-expression of L-R pairs, generation of heatmap (interaction activity map), and tissue plotting with contour marking tissue boundary is publicly available on github https://github.com/BiomedicalMachineLearning/SkinSpatial.

### Single molecule droplet digital PCR

Frozen scrolls were adjacently sectioned (110-120 µm total thickness) from the same OCT block that was used for Visium and RNAscope assays. Three individual BCC tissues from the block were isolated into separate tubes and were snap-frozen with dry ice. Total RNA was extracted using RNeasy MinElute Cleanup kit (Qiagen) according to the manufacturer’s instructions. RNA integrity was determined by Agilent RNA 6000 Pico kit and concentration was measured by Qubit (Thermo Fisher). cDNA was synthesized using Superscript^™^ IV VILO^™^ master mix with ezDNase^™^ enzyme (Invitrogen). In parallel, a No-RT control using equal RNA input was also generated to confirm the absence of gDNA contamination.

The ddPCR was carried out on the QX200 platform (Bio-Rad) according to the manufacturer’s instructions. Each triplicate reaction contained 1x ddPCR SuperMix for Probes no dUTP (Bio-rad), 1x target primer/probe mix conjugated with FAM or HEX (PrimePCR assay, Bio-Rad), cDNA, and dH_2_O. The controls consisted of a reaction mixture containing dH_2_O instead of cDNA or No-RT template from cDNA synthesis. Greater than 10,000 droplets were generated in each well by an automated droplet generator (range = 10,381 – 19,788). Subsequently, PCR amplification was performed in a C1000 Touch Thermal Cycler using an optimized program. The reaction was run at 95°C for 10 minutes, 40 cycles of 94°C for 30 seconds, 57.5°C for 30 seconds, and a final incubation at 98°C for 10 minutes. Results from the amplification were read using a QX200 Droplet Reader followed by data analysis with the QuantaSoft analysis software. The absolute transcript number for each target gene was determined by the software after manually setting the threshold for defining positive droplets. The mean of each triplicate was then calculated to give detected transcripts per microliter, from which values for transcript copies/ng RNA input were calculated.

### Generation of Vectra^®^ Polaris^™^ protein

Multispectral analysis of FFPE tissue utilised the MOTiF^™^ PD-1/PD-L1 kit (Akoya Biosciences, cat# OP-000001). Staining with Leica BOND RX (Leica Biosystems) and imaging with Vectra® Polaris^™^ (Akoya Biosciences) was performed at The Walter and Eliza Hall Institute (WEHI) Histology core facility as per kit manufacturer instructions. Briefly, tissue was stained through cycles of incubation with primary antibody, anti-IgG polymer HRP and covalent labelling with Opal TSA fluorophores, followed by heat induced epitope retrieval to remove bound antibodies prior to subsequent antibody cycles. Target markers included CD8 (Opal 480), PD-L1 (Opal 520), PD-1 (Opal 620), FoxP3 (Opal 570), CD68 (Opal 780, PanCK (Opal 690) and spectral DAPI DNA stain. Whole slide multispectral scanning was performed on the Vectra® Polaris^™^ using automatically adjusted exposure settings. Image tiles were spectrally unmixed in InForm® (Akoya Biosciences), then restitched in QuPath software (Bankhead et al., 2017).

### Applying STRISH for analysis of Vectra® Polaris^™^ mutiplex protein data

In addition to the analyses at transcriptomic level (RNAscope data), we also extended STRISH’s applications to quantify the interaction between cells at protein level. We reasoned that the STRISH pipeline is computationally flexible and could be applied to construct the landscape of L-R interaction at protein level. Similar to the analysis on RNAscope, STRISH first performed cell detection by applying positive cell detection on the image that was generated from the Vectra® Polaris^™^ system. Subsequently, the pipeline applied the PD-1 and PD-L1 markers detection and thresholding to the windows containing fewer than 100 cells. Finally, the STRISH min-max normalization was performed and a heatmap was plotted to display the local co-expression levels of PD-1 and PD-L1.

## Data and code availability

All the sequencing data and the H&E images used in this paper are publicly available in zenodo record (https://zenodo.org/record). The analysis script for STRISH cell detection and quantifying cell local co-expression is available at https://github.com/BiomedicalMachineLearning/SkinSpatial

