## SupplementaryFigures for "Spatial analysis of ligand-receptor interaction in skin cancer at genome-wide and single-cell resolution"

### Supplemental Figures:

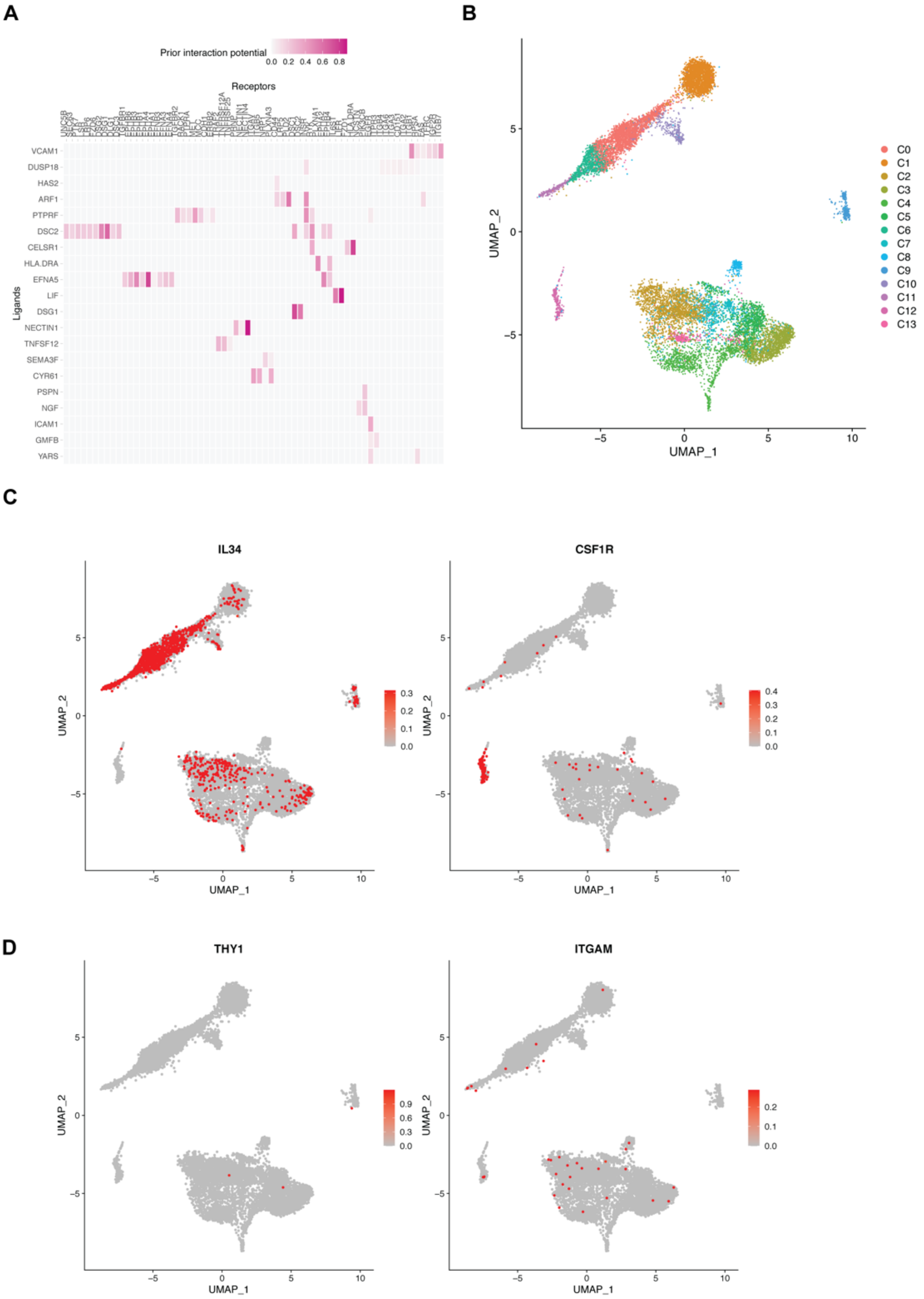


Supplemental Fig. S1. **scRNA-seq ligand-receptor interaction analysis for SCC cancer patient samples. (A)** NicheNet analysis results, with a heatmap showing ligand-receptor predicted interaction potential for the top 20 ligands. **(B)** A UMAP plot with cells grouped into 14 clusters using Louvain algorithm and gene expression. Each cluster is shown as one color. One dot represents a cell. **(C)** A feature plot highlighting (red dots) the distribution of cells expressing IL34 (left plot) and CSF1R (right plot). **(D)** A feature plot highlighting the distribution of cells expressing THY1 and ITGAM. Cells without expression appear as grey dots.

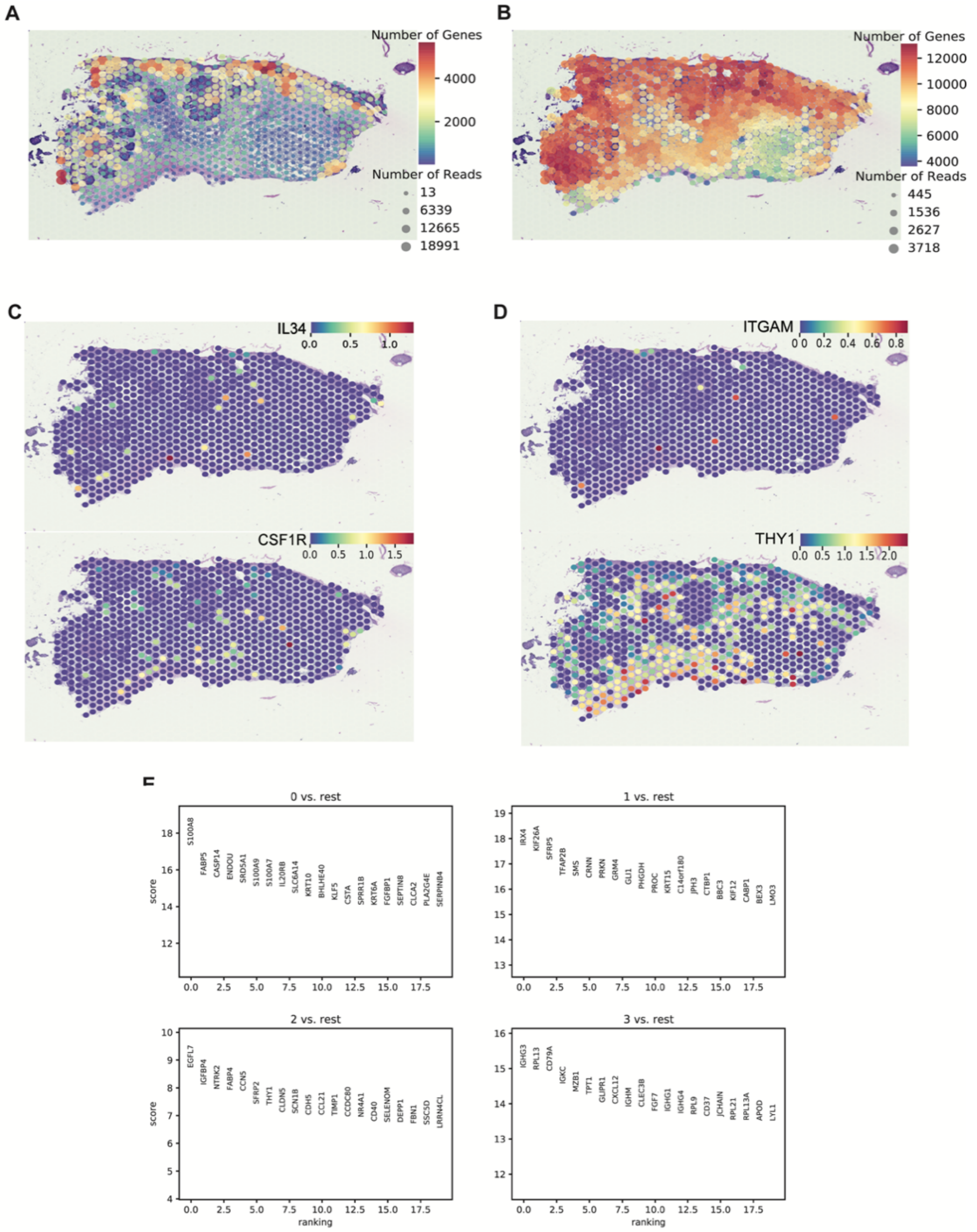


Supplemental Fig. S2. **ST-seq gene expression and cell-type analysis of the BCC skin cancer data.** **(A)** A QC plot showing the number of genes and reads captured in each spot across the whole BCC tissue section. **(B)** The number of transcripts captured in each spot after stLearn normalization. stLearn was applied to impute the signals of lowly expressed genes, using tissue morphological correlation to rescue zero values due to dropout (Pham et al, 2020). **(C)** The gene expression levels of IL34 and CSF1R (before normalization) displayed across all spots within the tissue. **(D)** A spatial feature plot showing the gene expression levels of ITGAM and THY1. **(E)** Analysis of gene markers for each of the four clusters identified from BCC Visium data (as shown in Figure 1). Each panel shows the top 20 genes most differentially expressed in one cluster compared to all the remaining clusters. The differentially expressed genes are odered from left to right of the X-axis, as from the most significant to the less significant (after Wilcoxon signed-rank test).

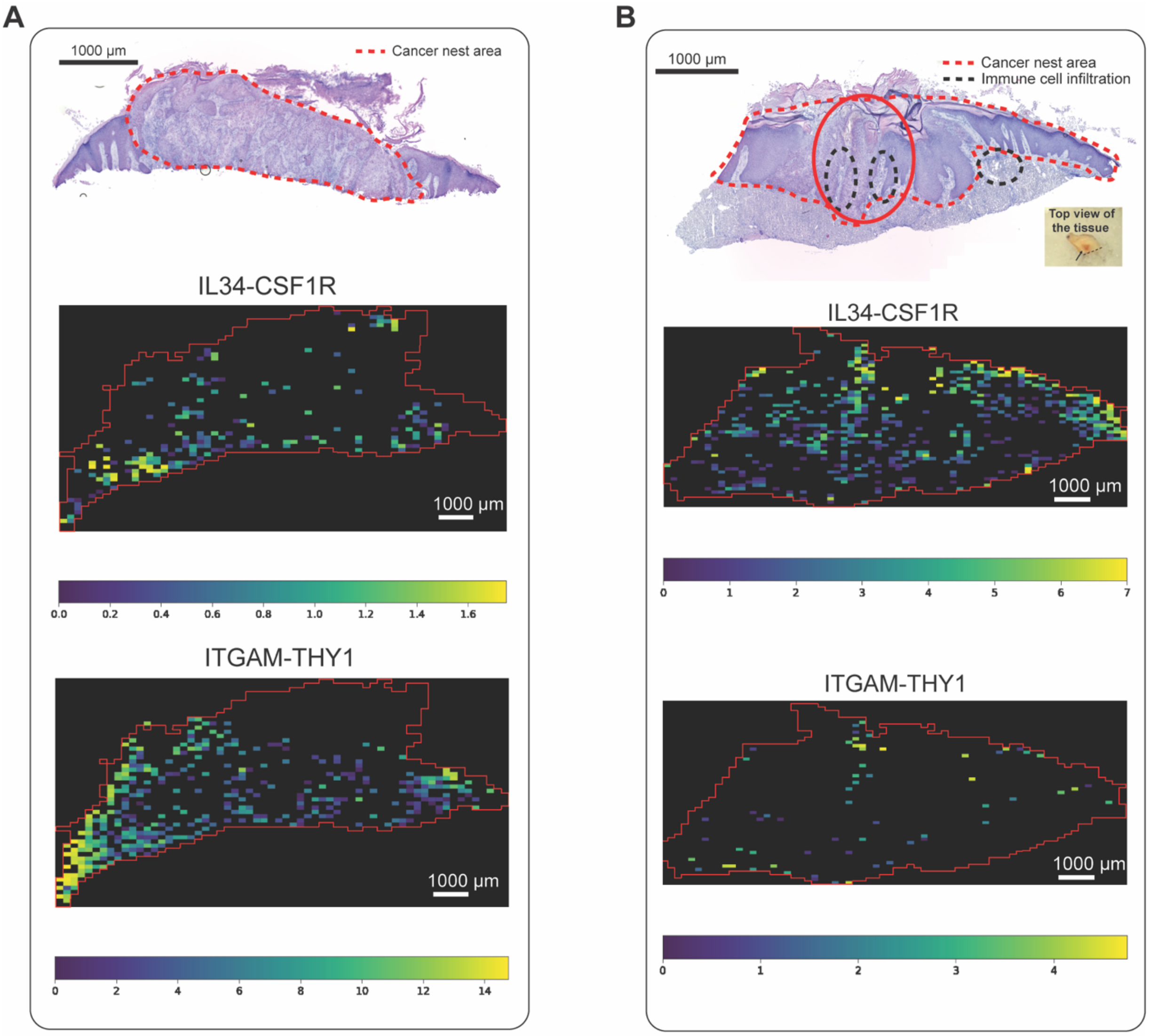


Supplemental Fig. S3. **Histopathological annotation and STRISH analysis of cell-to-cell interaction using RNAscope data of the two SCC cancer patient samples**. **(A)** From top to bottom, an annotated H&E image of a SCC patient ID-E15 and STRISH heatmaps of local co-expression for the two L-R pairs, IL34-CSF1R and THY1-ITGAM, respectively. **(B)** The top image is the annotated H&E of a SCC patient ID-F21. The red circle area indicates the same lesion core that indicates a black arrow in a small image at the bottom right corner. The dotted black line is indicating the side where the lesion tissue is embedded upright onto the OCT mold. The middle and bottom images are the heatmaps of the STRISH analysis of local co-expression of IL34-CS1R and THY1-ITGAM, respectively.

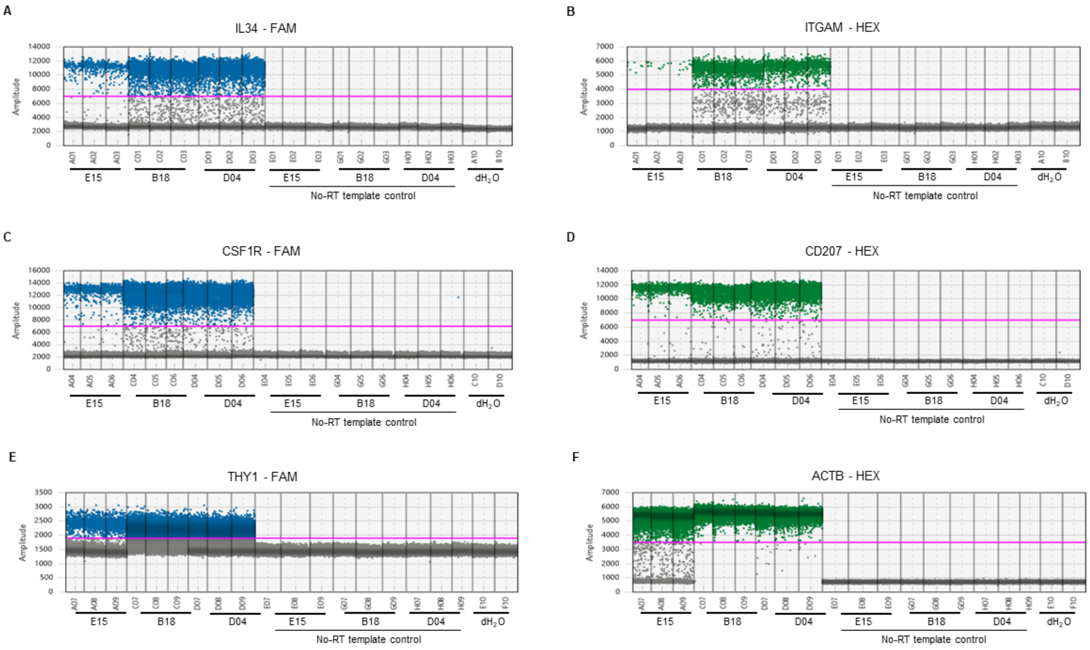


Supplemental Fig. S4. **Analysis of target genes by automated droplet digital PCR**. Expression plots for data from the three BCC patients and negative controls are shown. Each blue or green point represents a single droplet. The negative droplets were scored as grey if the fluorescent amplitude was lower than a detection threshold. The solid pink line indicates the threshold.

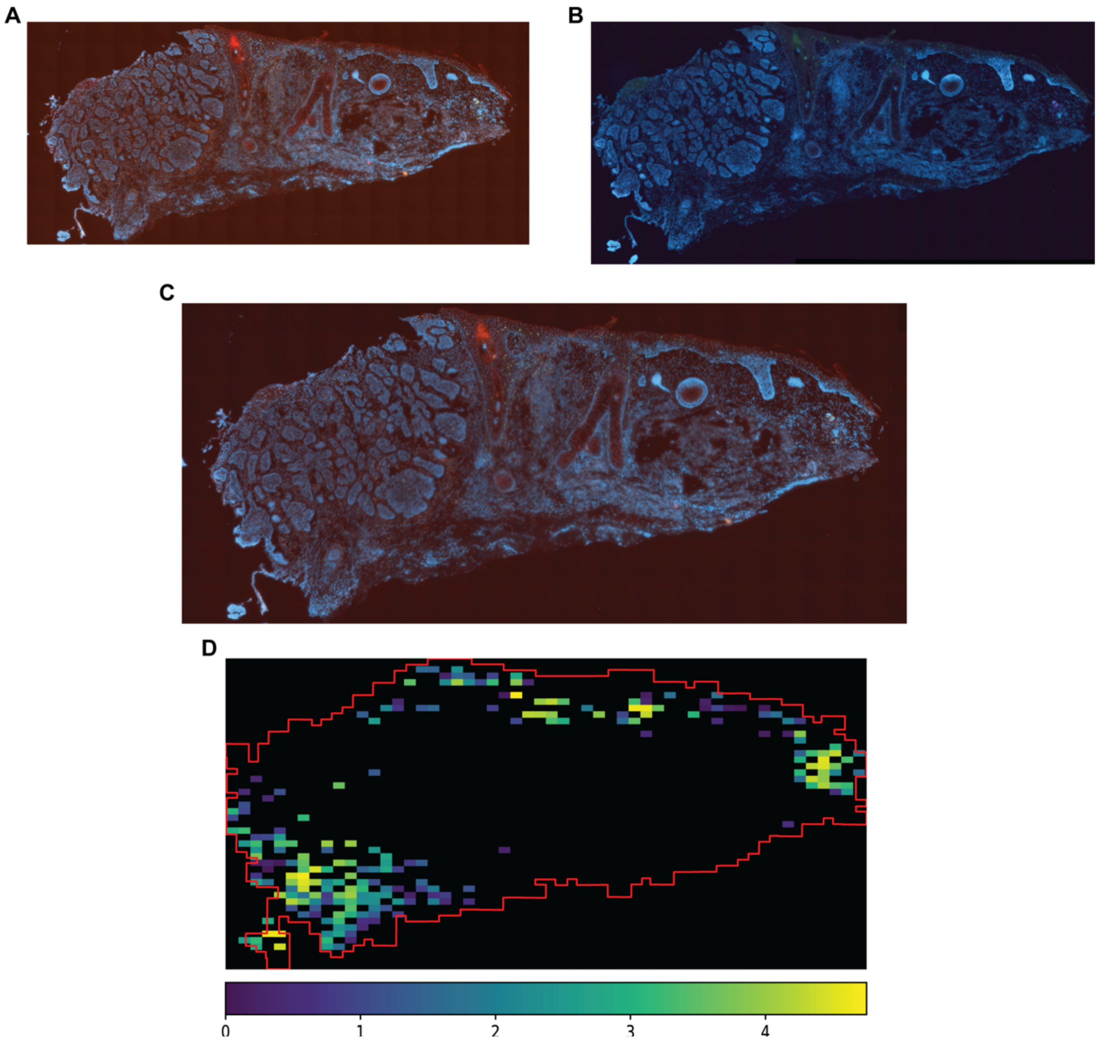


Supplemental Fig. S5. **Outputs from the STRISH analysis pipeline for RNAscope data. (A)** The RNAscope image for THY1, IL34, and CSF1R signals from the first round of imaging. **(B)** The RNAscope image for ITGAM and CD207 signals from the second round of imaging (for the same tissue section). **(C)** The result of image registration to combine the outputs from the two imaging rounds. After registration, the results from the two hybridization rounds (e.g. THY1 and ITGAM) were compatible with the local co-expression analysis. **(D)** STRISH analysis result showing the local co-expression of ITGAM and THY1 after image registration.
